# Inception: Simulating Personalized Long-Term Recovery in Disorders of Consciousness using Whole-Brain Computational Perturbations

**DOI:** 10.1101/2025.07.23.666344

**Authors:** Irene Acero-Pousa, Yonatan S. Perl, Jakub Vohryzek, Elvira G-Guzmán, Anira Escrichs, Ivan Mindlin, Jacobo Diego Sitt, Morten L. Kringelbach, Gustavo Deco

## Abstract

Advancements in the treatment of Disorders of Consciousness have seen significant progress with perturbative techniques and pharmacological therapies. Despite their potential, the underlying mechanisms of their variable efficacy remain poorly understood. To address this challenge, recent studies have utilised whole-brain modelling to simulate *in silico* perturbations. However, existing models focus exclusively on system behaviour during active stimulations, leaving unexplored how the brain dynamics evolve in post-acute and long-term stages, crucial in the recovery of consciousness. Here, we introduce *Inception*, a novel personalized approach to *in silico* perturbation modelling. We use a whole-brain models to simulate the perturbations and to unravel information about the long term effects of the proposed intervention. Applied to fMRI data from patients in a minimally conscious state and an unresponsive wakefulness state, our approach effectively simulates the transition to a healthy state, generating perturbed data closely resembling healthy brain activity. Moreover, we show that *Inception* enhances patient classification through machine learning, outperforming functional connectivity-based approaches. Finally, we investigate the correlation between perturbation responses and brain neuroreceptors, proposing that *Inception* might capture the long-term effects of pharmacological interventions in Disorders of Consciousness treatment.

## 1 Introduction

Following Crick and Koch’s foundational work on identifying the neural correlates of consciousness [1], the field has seen the development of numerous theories and definitions [2, 3], contributing to the ongoing debate around conscious experience. Although consciousness is inherently multidimensional [4], in clinical contexts it is characterized by two primary components: the content, which refers to awareness of oneself and the surrounding environment, and the level, associated with wakefulness [5]. Alterations in these components, whether induced by transient anaesthesia [6], brain injury [7], or intoxication [8], can result in profoundly altered states of consciousness.

A prominent example of such alterations is observed in Disorders of Consciousness (DoC), characterized by prolonged impairments in awareness and/or wakefulness. behavioural assessments such as the Coma Recovery Scale-Revised [9] are commonly used for diagnosis; however, they can misclassify up to 40% of conscious patients as unconscious [10, 11]. To improve diagnostic precision, quantitative tools such as fMRI, EEG low frequency power and EEG complexity indices have been proposed [12–14]. Nevertheless, the complex, dynamic, and multidimensional nature of consciousness continues to pose a significant challenge for accurate assessment. This complexity is particularly evident in the classification of minimally conscious state (MCS) and unresponsive wakefulness state (UWS) patients, due to the heterogeneous nature of the etiology, fluctuating arousal levels or ambiguous habituating responses [15]. MCS patients exhibit varying levels of awareness and responsiveness, ranging from minimal but definite behavioural evidence of self or environmental awareness (MCS+) to non-reflexive but reproducible behaviours (MCS-) [16]. In contrast, UWS patients show wakefulness without any discernible signs of awareness [17]. This variability underscores the critical need for personalized diagnostic and treatment approaches tailored to the unique profiles of each DoC patient.

The last decade has been crucial for advancing the understanding of DoC and improving patient recovery. Current treatment approaches include pharmacological interventions [18] and brain stimulation techniques [19]. Pharmacological interventions, such as zoldipem or amantadine [20], exert global effects on the brain by targeting neurotransmitter systems, particularly involving dopaminergic [18, 21] and noradrenergic [22] pathways. In contrast, brain stimulation techniques deliver invasive or non-invasive perturbations to localized brain regions. Examples include deep brain stimulation (DBS) [23], transcranial magnetic stimulation (TMS) [24] or transcranial direct current stimulation (tDCS) [25]. While some patients have shown promising responses to these treatments, their efficacy remains debated due to variability in their outcomes and the preliminary heterogeneous nature of existing evidence [7, 25]. More recently, research on pharmacological interventions has suggested the use of psychedelic substances, such as psilocybin, as a potential new treatment for DoC. The extensive literature on the effects of psychedelics, particularly on their agonism of the serotonin 2A (5-HT2A) receptor, indicates they may have the ability to elevate conscious awareness [26].

Although treatments for DoC have shown some success in improving patient outcomes, their implementation often relies on neuroanatomical hypotheses and clinical experience rather than a well-defined understanding of the underlying mechanisms, which remain poorly understood [27]. This gap in mechanistic insight limits our ability to predict when and why interventions will be effective. To address these limitations, recent efforts have turned to whole-brain modeling as a promising *in silico* approach for simulating brain stimulation [28–35]. These computational models integrate anatomical connectivity and functional dynamics obtained from neuroimaging data to reproduce brain activity. They allow for extensive experimentation with perturbations that model pharmacological or brain stimulation techniques, and circumvent the ethical concerns associated with empirical neurostimulation [36–38].

Such *in silico* paradigms have been instrumental in identifying causal mechanisms of recovery and determining perturbation parameters across neurodegenerative and neuropsychiatric disorders [28–33]. However, they focus exclusively on system behaviour during active perturbation, addressing only the initial stage of a broader framework. The subsequent post-acute effects, emerging after stimulation ends, provide valuable insights into the transient recovery dynamics of brain networks and how neural activity stabilizes, reorganizes, or decays in response to the intervention. Equally important are the long-term effects, which encompass the enduring changes in brain dynamics that arise from repeated or sustained perturbations [39]. Indeed, the recovery of DoC patients often manifests gradually over extended periods, typically spanning several months [40, 41], making post-acute and long-term outcomes particularly relevant.

To address this gap, we introduce *Inception*, a novel personalized approach to *in silico* perturbation-based whole-brain modeling designed to capture the post-acute and long-term effects of the perturbations. While existing models typically simulate the immediate dynamics generated by stimulating a single whole-brain model, the main innovation of *Inception* lies in the incorporation of a second model to simulate the downstream effects of these perturbed dynamics. This dual-model framework enables the simulation of brain dynamics as they transition toward a healthy state, even in the absence of continuous active perturbations. Specifically, *Inception* captures the long-term effect of external perturbations by embedding the induced changes within the brain’s underlying effective connectivity (EC). This allows to work in the generative space, enabling to observe how the changes propagate through the brain network and how they affect the brain dynamics over time.

We applied our methodology to fMRI data from two groups of DoC patients, namely MCS and UWS, to simulate their recovery. Our first hypothesis is that *Inception* effectively facilitates transitions toward a healthy brain state and thereby capturing the long-term effects of perturbations. Once the healthy state would be achieved, we would seek to identify the network characteristics of the brain regions most conducive to recovery after stimulation. Moreover, we hypothesize that the subject-specific perturbations needed for recovery would be informative for improving diagnostic criteria when integrated with machine learning models. On the other hand, we hypothesize that the brain regions more susceptible to recovery would share a neuroreceptor profile [42], contributing to the understanding of the long-term effects of pharmacological interventions in DoC. Lastly, we hypothesize that *Inception* outperforms traditional *in silico* methods in inducing a whole-brain recovery state.

## 2 Results

The *Inception* framework consists of three steps, each generating three output matrices: effective connectivity (EC), functional connectivity (FC), and lagged-covariance (COV). EC reflects the directional influences between brain regions that sustain specific brain dynamics over time and is inferred through a whole-brain model. FC captures the pairwise correlations between brain regions, indicating the degree of coordination between them, and can be derived from either empirical data or model simulations. Similarly, COV represents the time-lagged covariance between regions, providing insights into causal relationships, and can also be computed from empirical data or simulated with a model.

The framework begins by defining the initial and target brain states (Figure 1b). The initial state represents the brain dynamics of each UWS and MCS patient, simulated using a whole-brain Hopf model (see Methods) to derive their **individual EC, FC**, and **COV**. The target state corresponds to the dynamics of a healthy group, obtained by simulating brain activity for each healthy subject, computing their individual EC, and averaging across individuals to derive a representative **target EC**.

**Figure 1.**
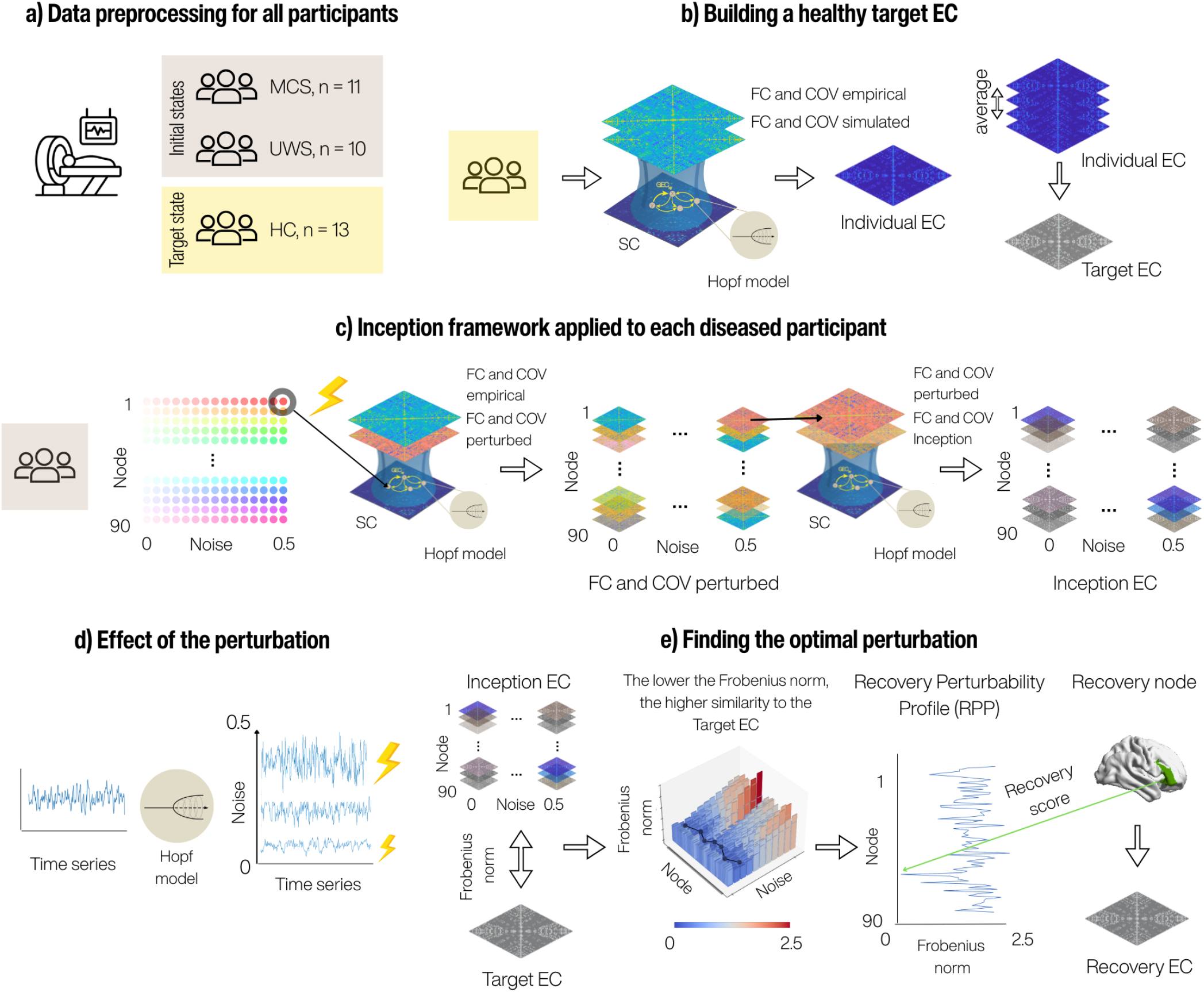
Overview of the data and the *Inception* framework. **a** We analyzed fMRI data from 21 patients diagnosed with two Disorders of Consciousness: Minimally Conscious State (MCS, n = 11) and Unresponsive Wakefulness Syndrome (UWS, n = 10), along with 13 Healthy Controls (HC). The goal is to perturb the initial states (MCS and UWS) to induce dynamics resembling the target healthy state (HC). **b** The target Effective Connectivity (EC) represents the desired connectivity profile. It is constructed by fitting whole-brain Hopf models to each HC subject’s data to generate their individual EC matrices, which are then averaged to form the group-level Target EC. **c** The *Inception* framework is applied individually to each MCS and UWS subject. Perturbations are introduced by systematically increasing the Gaussian noise parameter (*η*) at each brain node (range: 0 to 0.5) within the subject’s whole-brain model. This generates perturbed functional connectivity (FC), time-lagged covariance (COV), and EC matrices. These are then input into a second, non-perturbed model (with uniform noise across nodes), yielding the so-called Inception FC, COV, and EC, which capture both post-acute and long-term effects of the perturbations. **d** Perturbation effects are driven by modulating the noise intensity at individual nodes. Higher noise levels result in time series with increased amplitude. **e** For each generated Inception EC, similarity to the Target EC is assessed using the Frobenius norm. The minimum Frobenius norm across all noise levels at each node defines the Recovery Perturbability Profile (RPP). The node with the lowest norm (i.e., highest similarity to the target) is termed the recovery node, its value the recovery score, and the corresponding Inception EC is referred to as the recovery EC.

In the second step (Figure 1c), each initial brain state undergoes *in silico* perturbation. Perturbation is applied to a specific node by modifying the parameter of Gaussian noise, thereby modulating the node’s activity to mimic an external intervention. Within this newly defined parameter space, the model is optimized to generate three outputs: the **perturbed EC, FC** and **COV** matrices, capturing the effects of the active perturbation. This process is repeated systematically for all nodes and for perturbation intensity values ranging from 0 to 0.5.

The third and novel step of the *Inception* framework involves feeding the output matrices of the perturbed model into a subsequent model that operates without external perturbation. This step simulates the evolution of the perturbed brain state to capture the long-range effects of stimulation. Through this process, the model generates an EC matrix that encapsulates these effects, referred to as the **Inception EC**, alongside the corresponding **Inception FC** and **Inception COV**.

To assess the similarity of the perturbed system to the target healthy group (i.e., recovery), we developed the **Recovery Perturbability Profile (RPP)** (Figure 1d). The RPP evaluates the potential of each brain node to facilitate recovery by quantifying its ability to drive the system toward the target state. Specifically, for each node, we computed the Frobenius norm distance between the Inception EC obtained at varying perturbation intensities and the target EC. From this range of a given node, we identified the perturbation intensity that minimized the distance and included this distance value as its score in the RPP. The smallest value in the RPP, referred to as the **recovery score**, is associated with the node with the highest potential to achieve similarity to the target EC, and termed the **recovery node**. Finally, to highlight the Inception EC, FC and COV derived from stimulating this recovery node at its optimal perturbation intensity we name them **recovery EC, FC and COV** representing the closest approximation of the perturbed system to the healthy target state.

### 2.1 *Inception* effectively facilitates transitions toward a healthy brain state

First, we aimed to evaluate the effectiveness of the perturbations in achieving a successful transition to the target healthy state (Figure 2). To do this, we compared the individual EC of each HC and DoC patients to the healthy target EC, as well as the recovery EC (R) of each DoC subject to the healthy target EC, using the normalized Frobenius norm as a measure of similarity.

**Figure 2.**
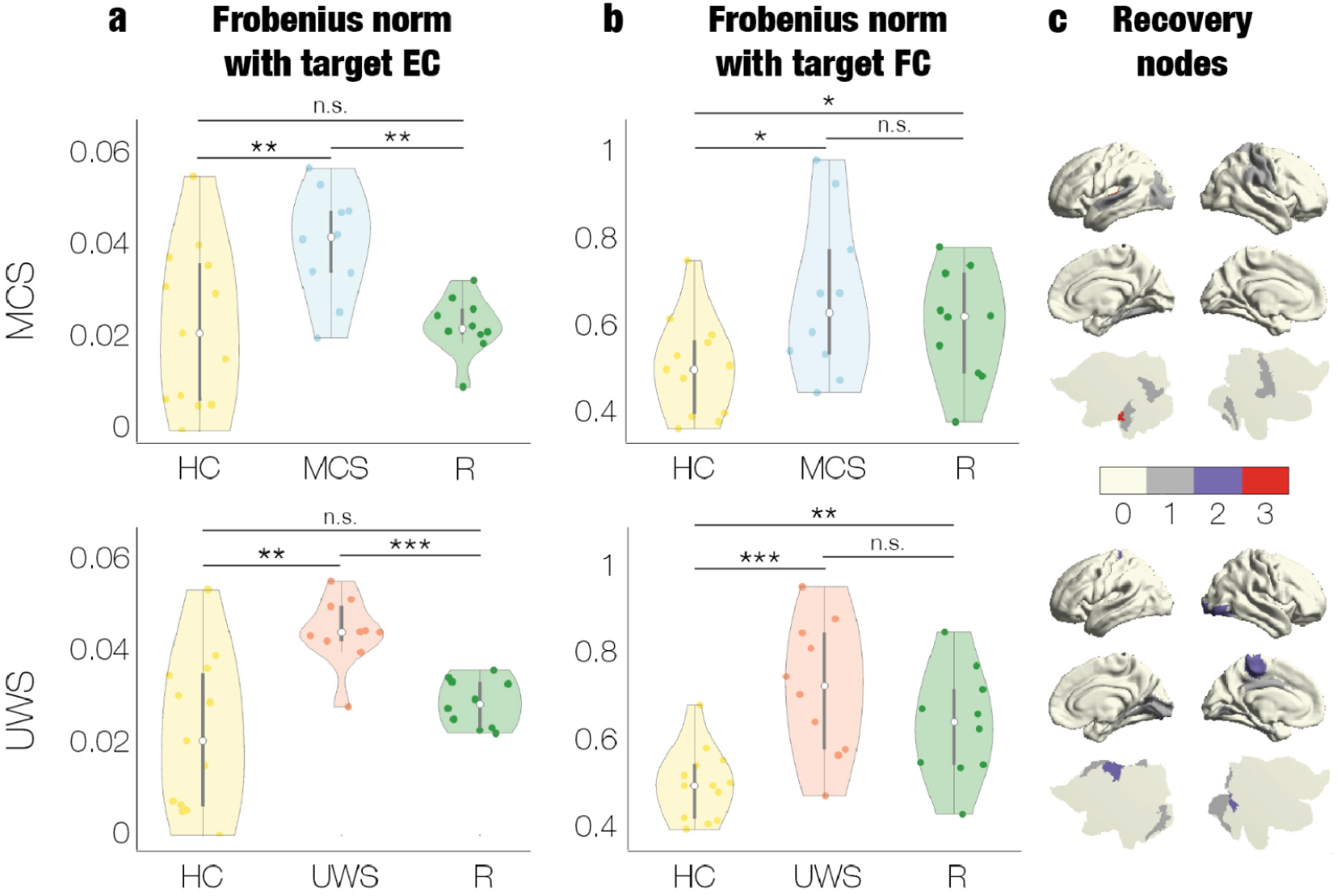
Node-specific perturbations drive *in silico* recovery towards the healthy state. **(a)** Similarity between the Target EC (computed as the average EC of healthy controls, HC) and the individual ECs of HC, minimally conscious state (MCS; top), unresponsive wakefulness state (UWS; bottom) subjects, and the recovered EC (R), measured using the normalized Frobenius norm. **(b)** Similarity between the Target FC (computed as the average simulated individual FC of HC) and the simulated individual FCs of HC, MCS (top), UWS (bottom) subjects, and R, also measured using the normalized Frobenius norm. Significance levels are indicated as follows: * p *<* 0.05, ** p *<* 0.01, *** p *<* 0.001. **(c)** Brain renderings of recovery nodes identified in the MCS (top) and UWS (bottom) groups. The color scale indicates the number of subjects for whom each node was identified as a recovery node.

As shown in Figure 2a, for the MCS group (top), the similarities of the HC group were not significantly different (p *>* 0.05) from those in R, indicating an optimal transition to the healthy target state. Moreover, the similarity was significantly different when comparing the MCS with both HC and R (HC vs. MCS, p *<* 0.01, R vs. MCS, p *<* 0.01). On the other hand, the UWS group (bottom) also achieved an optimal transition, with the similarities between HC and R not showing significant differences (p *>* 0.05), but both R and HC being statistically different from the UWS group (p *<* 0.05).

Furthermore, we investigated if the recovery was also present in the functional domain (Figure 2b). The functional domain is assessed using the target FC, which we computed as the average simulated FC of all HC. We followed the same procedure, focusing on the normalized Frobenius norm as an index of similarity. In this case, we compared the simulated individual FC of each HC and DoC patient and their recovery FC (R) to the target FC. Although the comparison between the HC and R is significant, there is a clear trend indicating partial recovery both for MCS and UWS.

We next focused on distinguishing the recovery nodes in each group. Figure 2c illustrates the frequency with which each node was identified as the recovery node across subjects. For MCS (Figure 2c, top), the left heschl gyrus was the most effective node, identified as optimal in three out of eleven subjects, followed by the middle occipital gyrus, left the left caudate nucleus, the left putamen nucleus, the left superior temporal gyrus, the right supramarginal gyrus, the right postcentral gyrus, the right fusiform gyrus and the right hippocampus, each found to be optimal once. For the UWS (Figure 2c, bottom), the left paracentral lobule and the right inferior occipital gyrus were the most effective nodes, appearing as optimal in two out of ten subjects each. This was followed by the median cingulate and paracingulate gyri, the left parahippocampal gyrus, the left lingual gyrus, the right lingual gyrus, the right calcarine fissure and the right amygdala, each observed as optimal once.

### 2.2 Network characteristics of nodes supporting recovery

We wanted to understand the nature of nodes that are most effectively perturbed to achieve the target healthy state. For that, we correlated each node’s score in the RPP with its EC degree. This analysis was conducted both at group and individual levels. At the group level (Figure 3a), we examined the mean RPP across all subjects in each group and its correlation with the corresponding group-level EC degree. At the individual level (Figure 3b), we correlated each subject’s RPP with its individual EC degree. We found significant positive correlations in both the MCS (r = 0.27, p *<* 0.01) and UWS (r = 0.39, p *<* 0.001) groups. Significant correlations were present in all subjects except one in the MCS group and all but two subjects in the UWS group. These results suggest that nodes with lower EC degrees are more effectively perturbed and more capable of recovering toward the healthy target state.

**Figure 3.**
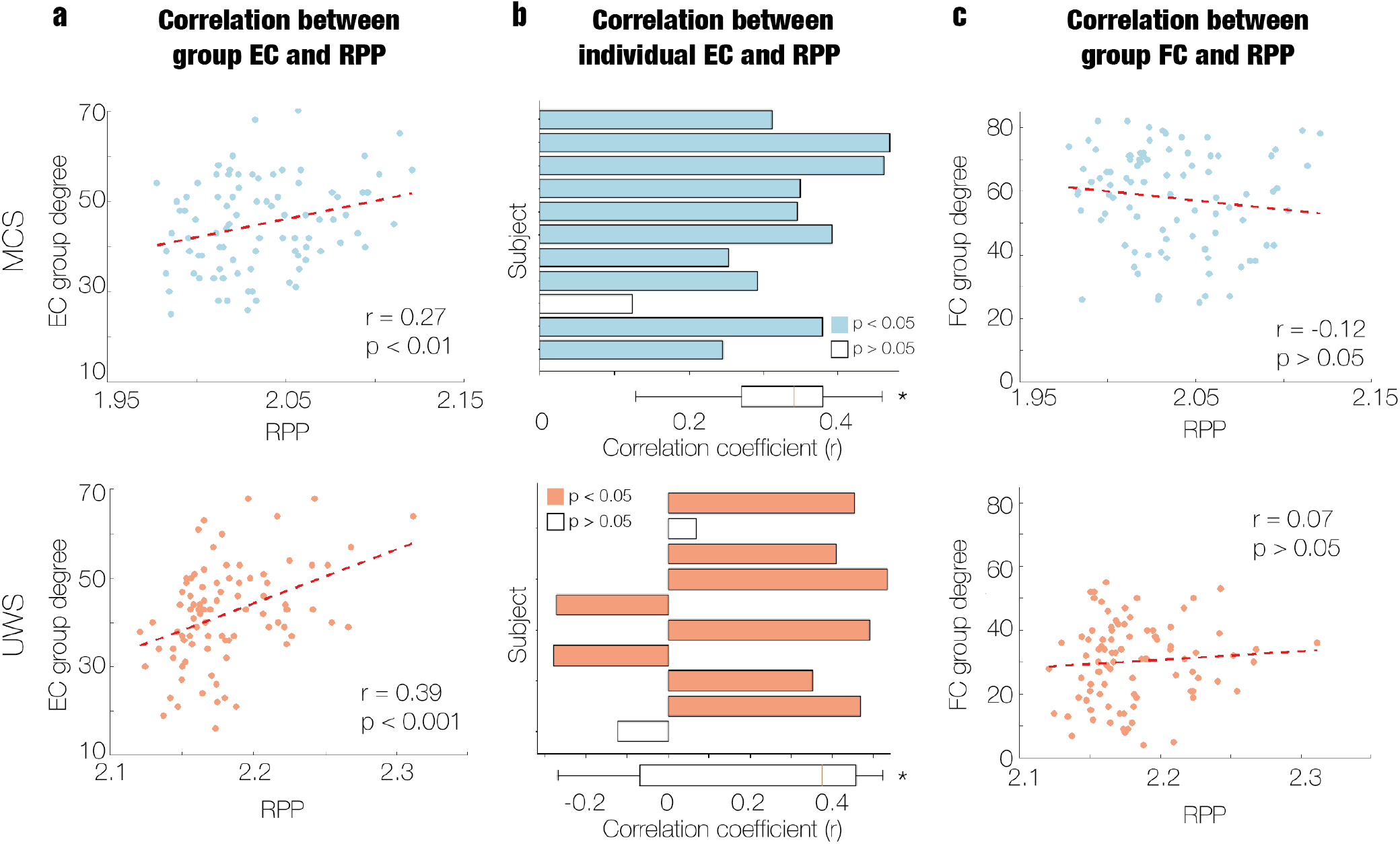
Lower EC degree characterizes nodes supporting recovery. **(a)** Correlation between the RPP and the degree of EC at the group level for MCS (top left) and UWS (bottom left) groups. **(b)** Correlation between the RPP and the degree of EC at the individual subject level for MCS (top middle) and UWS (bottom middle) subjects. Bars represent individual subjects, with significant correlations (p *<* 0.05) indicated in color. The boxplots show the distribution of correlation values across subjects in each group, providing a statistical comparison against a non-correlation baseline. **(c)** Correlation between the RPP and the degree of the empirical FC at the group level for MCS (top right) and UWS (bottom right) subjects.

We also explored whether the observed results extended to the functional connectivity domain at the group level. Using the group empirical FC, we calculated the degree by applying a threshold to the correlation values, where any correlation below a specified cutoff (ranging from 0 to 0.5) was set to zero. However, no significant correlations were observed across all tested thresholds (p *>* 0.05). For illustration, we present the correlation at a threshold of 0.25 in Figure 3c. As shown, there was no significant relationship between the group RPP and the degree of group-level FC (MCS: r = −0.12, p *>* 0.05; UWS: r = 0.07, p *>* 0.05). These results suggest that EC, rather than FC, plays a more critical role in capturing the effects following a perturbation.

### 2.3 Subject-specific perturbations leading recovery enhances diagnostic precision of DoC

To investigate the diagnostic potential of the RPP in distinguishing between the two DoC groups, we compared the performance of 1000-fold k-means clustering using individual RPP, and individual EC and FC matrices from MCS and UWS patients (Figure 4a). To ensure consistent dimensionality across the inputs for each clustering analysis, we computed the global brain connectivity (GBC) from the FC matrices and the degree of the EC matrices. This step resulted in input vectors of size 1×90 for all data types, matching the dimensionality of the RPP.

**Figure 4.**
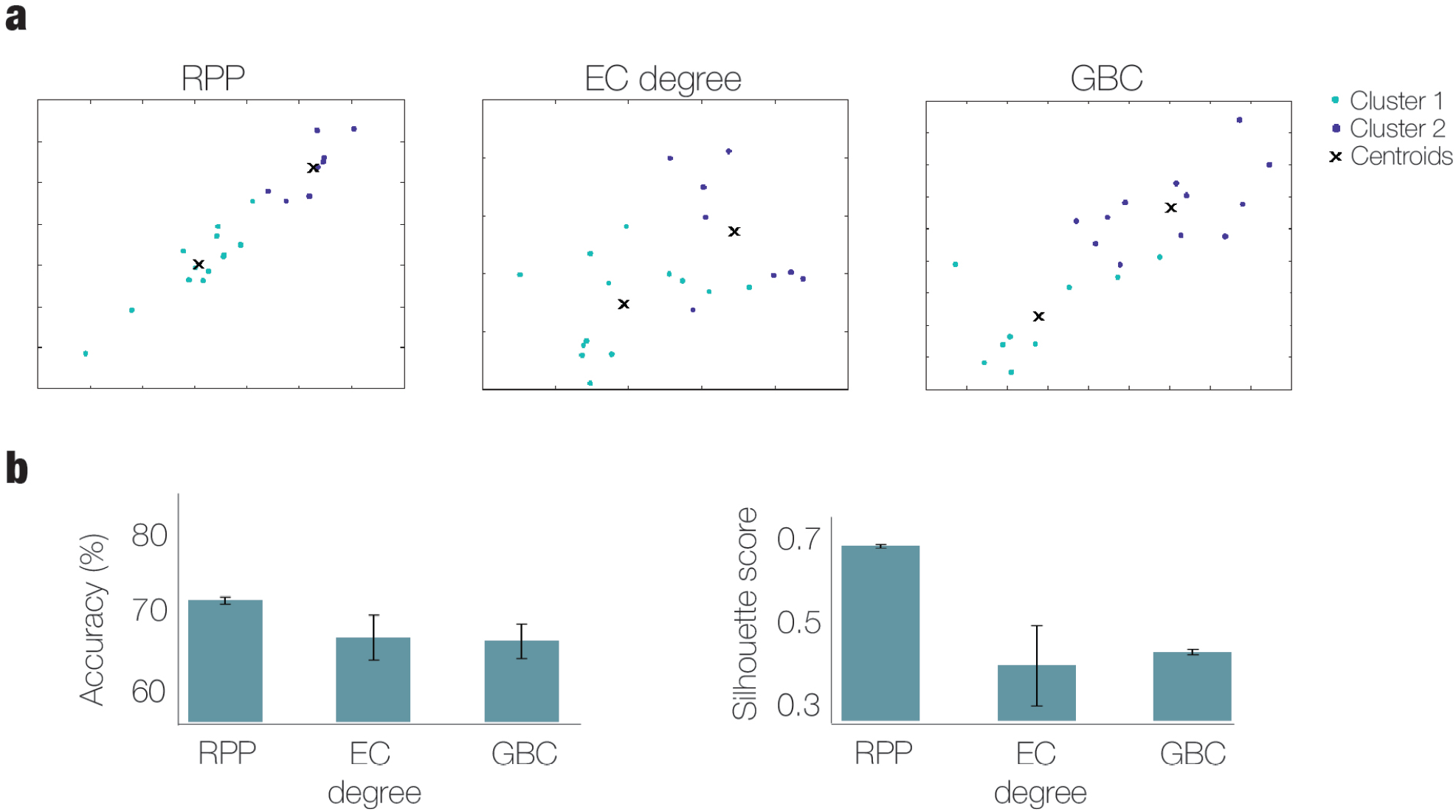
Node-specific recovery (RPP) enhances the diagnostic precision. **(a)** Results of the 1000-fold k-means clustering analysis applied to individual RPP, EC degree, and GBC measures for MCS and UWS subjects. The EC degree was computed from the individual EC matrices, and the GBC from the simulated individual FC matrices. The scatter plots display the clustering results, with clusters denoted by different colors (Cluster 1 in cyan, Cluster 2 in purple) and centroids are marked by black crosses. **(b)** Performance evaluation of clustering using accuracy (left) and silhouette score (right) metrics. The black bar indicates the variance obtained across the 1000-fold.

Clustering performance was evaluated using accuracy and Silhouette score metrics (Figure 4b). RPP outperformed the other inputs, achieving an accuracy of 71.43% and a Silhouette score of 0.69. In contrast, EC total degree and GBC measures showed lower performance: EC resulted in an accuracy of 64.97% and a Silhouette score of 0.42, while GBC achieved an accuracy of 64.40% and a Silhouette score of 0.45.

### 2.4 Neuroreceptor composition of nodes leading to recovery

Next, we aimed to determine the neuroreceptor composition of the nodes associated with recovery, as computed by *Inception*. To do so, we examined the correlation between the averaged recovery potential (RPP) values per group and the distribution of serotonergic (5HT1a, 5HT1b, 5HT2a, 5HT4, 5HT6, 5HTT), dopaminergic (D1, D2, DAT), and norepinephrinergic (NET) receptors. The results of this analysis are shown in Figure 5.

**Figure 5.**
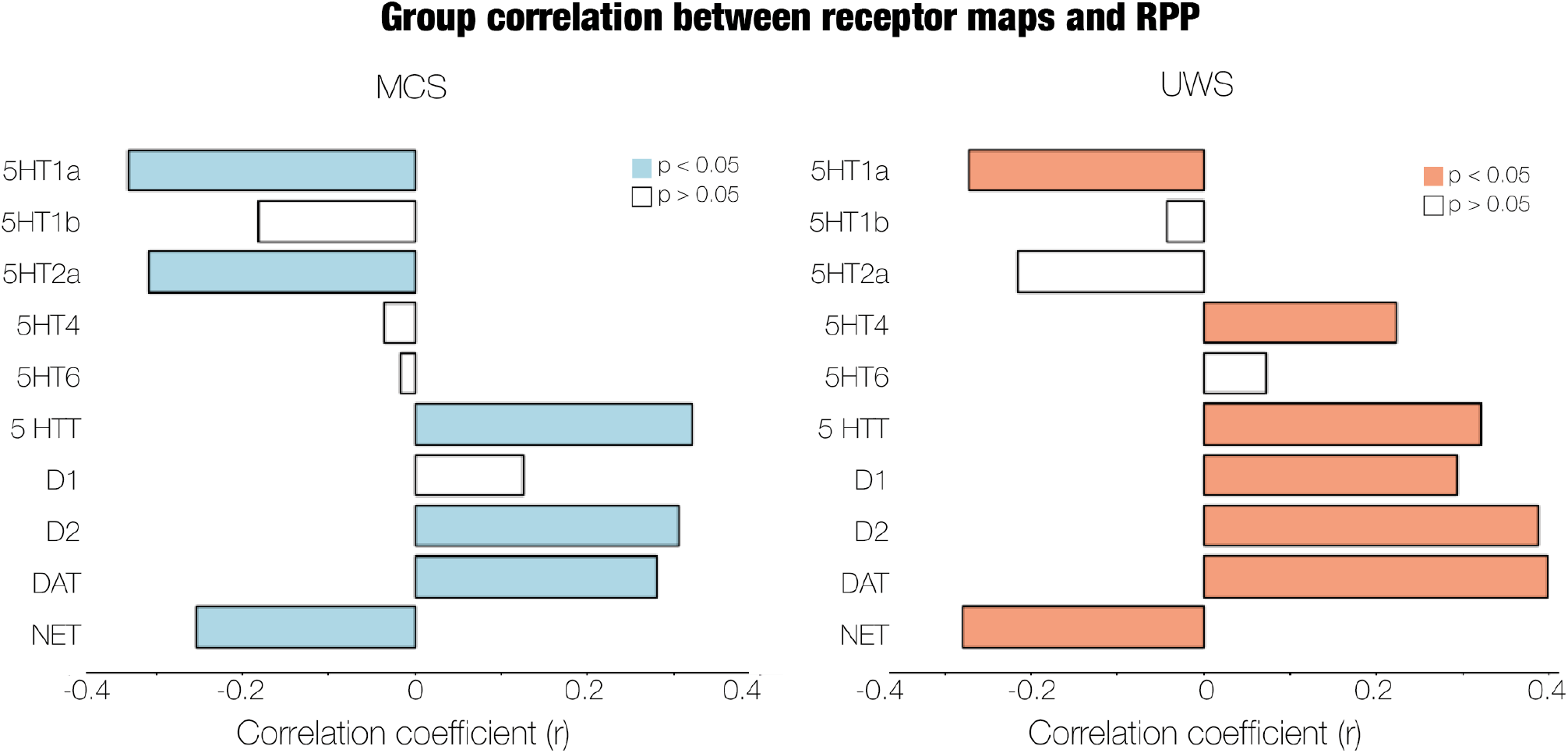
Node-specific recovery (RPP) correlates with receptor map distribution. Correlation between the RPP and receptor maps at the group level for MCS (left) and UWS (right) groups. Bars represent each receptor map, with significant correlations (p *<* 0.05) indicated in color.

In the MCS group (Figure 5, left), significant negative correlations (p *<* 0.05) were observed for serotonin receptors (5HT1a and 5HT2a) and norepinephrine transporters (NET), indicating that higher densities of these receptors are associated with enhanced recovery capacity. In contrast, significant positive correlations (p *<* 0.05) were identified for serotonin transporter (5HTT) and dopamine receptors (D2 and DAT). These results suggest that lower densities of these receptors within a node are linked to better recovery trajectories.

For the UWS group (Figure 5, right), we identified significant negative correlations (p *<* 0.05) for serotonin receptor 5HT1a and norepinephrine receptor NET, again implying that increased receptor density correlates with greater recovery potential. In contrast, significant positive correlations (p *<* 0.05) were found for serotonin receptor 5HT4, the serotonin transporter (5HTT), and dopamine receptors (D1, D2 and DAT). This indicates that lower receptor densities in these nodes are associated with improved recovery outcomes.

All correlations were tested against 1000 randomized versions of the receptor maps preserving spatial autocorrelations (see Supplementary 1) [43]. All were found to be significantly different from the null distributions.

### 2.5 Comparing *in silico* perturbation-based models: *Inception* and *Awakening* frameworks

Finally, we wanted to assess whether *Inception* provides more accurate identification of perturbations leading to a healthy state compared to previous traditional *in silico* frameworks. For this comparison, we selected *Awakening* (introduced by Deco et al. [28], see Methods), which is based assessing the simulated dynamics of a model incorporating active perturbations. In contrast, *Inception* includes a second model to account for the long-term effects of stimulation.

Using the perturbed EC we computed the RPP of *Awakening* following the same procedure as in *Inception*. We then compared the recovery scores, defined as the minimum value in the RPP, between the two approaches. Furthermore, we replicated the analysis from the FC and COV perspective, building their RPP, and finding their recovery score.

As illustrated in Figure 6, *Inception* outperformed *Awakening* across all metrics and groups, with significantly lower RPP values, indicating stronger similarity to the healthy target state. In the MCS group (Figure 6, top), we observed a significant lower distance for the EC analysis (left, p *<* 0.01), as well as for the FC (middle, p *<* 0.05) and COV (right, p *<* 0.05) comparisons. In the UWS group (Figure 6, bottom), *Inception* showed more pronounced improvements, with the EC comparison yielding p *<* 0.001, the FC comparison p *<* 0.01, and the COV comparison p *<* 0.05.

**Figure 6.**
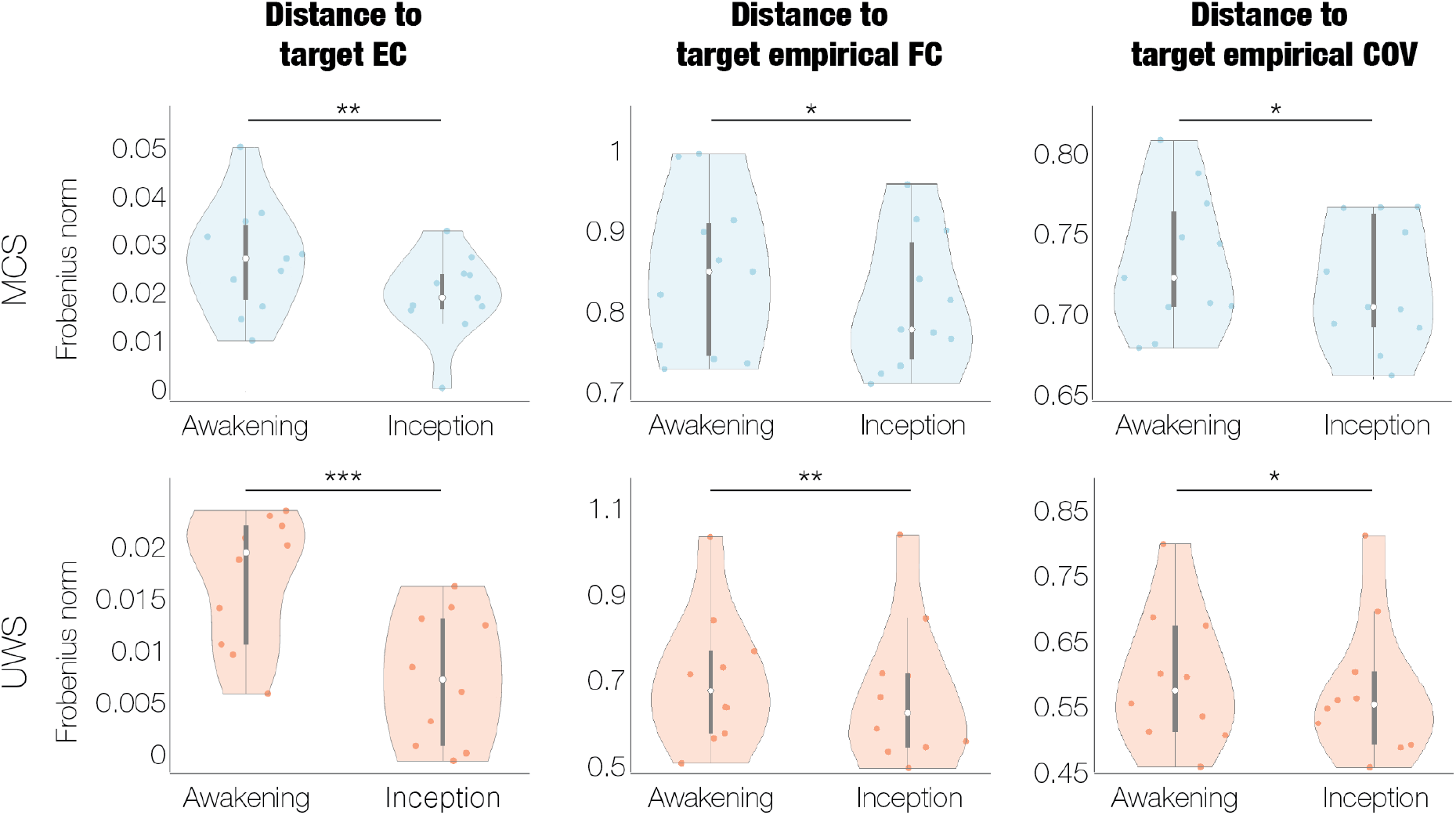
Comparison of *in silico* perturbation-based models: *Inception* and *Awakening* frameworks. Violin plots depicting the normalized Frobenius norm distances to target EC (left), FC (middle), and COV (right) matrices for each subject of MCS (top row) and UWS (bottom row) groups. In each plot, the distances achieved using the *Inception* framework are compared with those using the *Awakening* framework. Significance levels are indicated as follows: * p *<* 0.05, ** p *<* 0.01, *** p *<* 0.001.

## 3 Discussion

In this study, we introduced *Inception*, a novel personalised *in silico* perturbation-based whole-brain modelling approach for simulating post-acute and long-term effects of brain perturbations. This methodology accurately models the transition of individual’s brain dynamics from unresponsive wakefulness state (UWS) or minimally conscious state (MCS) to a healthy state. Our analysis identified the key recovery regions of each patient, demonstrating that perturbing nodes with lower effective connectivity (EC) degree yields more effective recovery outcomes. Moreover, we demonstrated that using the patient-specific recovery potential of brain regions (i.e., the Recovery Perturbability Profile (RPP)) in a k-means clustering model enhances individual classification accuracy compared to measures based on EC or FC. Additionally, we found significant correlations between the RPP and the distribution of serotonergic, dopaminergic and noradrenergic neurotransmitter systems [42]. Finally, we demonstrated that *Inception* outperforms traditional *in silico* perturbation-based models.

Whole-brain modelling has been extensively used to simulate and understand the response of brain dynamics to perturbations, opening possibilities for identifying novel targets for DBS or TMS in health and disease [28–34]. While valuable for identifying immediate responses to perturbations, prior approaches have neglected how brain dynamics evolve after the cessation of stimulation. As a result, the postacute and long-term effects of perturbations, which have proven critical for the recovery of consciousness [40, 41], remain largely unexplored. To address this gap, we developed *Inception*, extending traditional perturbation-based modelling by capturing long-term post-stimulation dynamics. To achieve this, we first used a whole-brain model to explore *in silico* perturbations, simulating active perturbation responses; and second, a subsequent unperturbed whole-brain model, simulating the evolution of these perturbed dynamics over time. Similar to the film *Inception*, where agents delve into dreams to uncover hidden states of the mind, our approach reveals the underlying mechanisms driving the long-term effects triggered by perturbations.

We used EC as the metric to embed the captured effects of the stimulation, differing from traditional methodologies that relied solely on FC. The reason behind this choice lies in the nature of the information captured by each metric. EC explains the causal influence that one neural system exerts over another, modelling the directed, context-dependent interactions between brain regions. In contrast, FC measures undirected statistical dependencies between brain regions, without distinguishing the direction of influence [44–47]. While FC can identify correlated patterns of activity, it does not provide insights into the underlying causal mechanisms driving those correlations. Being able to capture directional changes is essential in the context of modelling perturbations, as the stimulation of one brain region can significantly alter its impact on other regions. This makes EC more appropriate for embedding the effects of the perturbation. Furthermore, recent work confirms that the efficacy of non-invasive brain stimulation depends on white-matter structure, that is, anatomical connectivity [48–50]. In addition to these studies, [51] found that the structural connectivity explains local and global effects of brain stimulation. Thus, EC, which is built on the structural connectivity, offers a robust framework to integrate the post-acute and long-term effects of stimulations.

After applying *Inception* to each MCS and UWS patient, we identified the recovery nodes, that is, the ones achieving a better transition to the healthy state after a perturbation. In the MCS group, the left Heschl’s gyrus and the left superior temporal gyrus emerged as particularly significant recovery nodes, underscoring the crucial role of the auditory cortex. These findings align with existing research that highlights the potential of auditory stimulation as a non-invasive therapeutic approach for DoC patients, with particular emphasis on the effectiveness of calling a subject’s own name [52–54]. Also, it has been shown that a higher-level activation of the auditory cortex in response to a familiar voice speaking the patient’s name suggests an increased chance of recovering consciousness within 12 months [55]. Together with these regions, some subcortical areas were also found important for the transition to the healthy state, including the left caudate nucleus, the left putamen nucleus and the right hippocampus.

Interestingly, a previous study found that these three structures increased significantly in volume after a DBS implant that led to functional recovery, potentially suggesting their recovery potential in patients with DoC [56]. Previous research has observed that the emergence from the MCS is characterized by the recovery of functional interactive communication or functional use of two different objects [57]. Many of the regions found as recovery nodes support these two milestones, especially the supramarginal gyrus and superior temporal gyrus due to their role in language processing, and the fusiform gyrus, involved in object identification.

In the UWS group, two regions were found most commonly as recovery nodes, the left paracentral lobule and the right inferior occipital gyrus. This first region mentioned was found to be the most sensible area in MCS subjects when applying the *Awakening* framework in DoC patients [31]. It belongs to the primary motor cortex (M1), which has widely been observed to promote recovery of motor impairment when stimulated with TMS [58]. On the other hand, the occipital area, where the right inferior occipital gyrus and the right calcarine fissure are, has previously been found crucial for the recovery of UWS patients [59]. Together with these areas, the left and right lingual gyrus also belong to the visual cortex and were marked as recovery nodes. This finding is consistent with prior research, which states that visual orientation towards stimuli and visual tracking are crucial for recovery and, moreover, it has been stated that vision is important for the recovery as without it active manipulation of objects may be misguided or not initiated [59, 60].

We explored the factors underlying a node’s potential for recovery after perturbation. Our analysis indicates that regions with weaker EC degree are more likely to promote a successful transition to the healthy state. This finding goes in line with previous studies that suggested that the network periphery tends to exhibit greater variability, adaptability, and plasticity, whereas the network core primarily supports system robustness and stability [61–63]. Furthermore, Bassett et al. [64] demonstrated core stability and peripheral flexibility during a learning task using whole-brain fMRI data. Additionally, the higher correlation strength in the UWS group may reflect a heightened dependence on low-connectivity nodes for recovery, possibly due to more extensive network disruptions associated with this state [65]. Nonetheless, the overall moderate strength of these correlations indicates that EC degree is but one of several factors influencing recovery potential. Factors such as a node’s functional role, network location, neurochemical environment, and anatomical properties also impact to determine its responsiveness to perturbation [58, 66]. In contrast, we did not observe a significant correlation between recovery potential and functional connectivity, aligning with previous research that suggests the transmission of TMS is not strongly predicted by FC [67].

The complexity of diagnosing Disorders of Consciousness (DoC) and the variability in patients’ responses to treatment highlight the need for personalized treatment strategies. Reliable biomarkers are key to enhancing diagnostic accuracy and guiding interventions that align with each patient’s unique neural dynamics and recovery potential. Interestingly, our study demonstrates that the Recovery Perturbation Profiles (RPP) can be used for accurate diagnosis of DoC, significantly outperforming traditional EC and FC measures [68, 69]. This superior accuracy of RPP may stem from its ability to capture how the brain responds to perturbations, integrating both baseline connectivity and the network’s dynamic recovery potential. While FC measures static correlations between brain regions and EC examines causal interactions, RPP uncovers latent recovery processes and neural plasticity, establishing it as a more robust model-based biomarker for DoC diagnosis.

Pharmacological treatments for DoC are commonly used to modulate neurotransmitter systems and induce widespread changes in brain activity. Given that *Inception* models long-term perturbation effects, we investigated whether it could capture these pharmacologically relevant dynamics. Our results regarding serotonin align with previous studies demonstrating the restoration of brain dynamics indicative of wakefulness following the stimulation of serotonergic receptors [70]. Specifically, 5HT2A serotonin receptor has been proposed as a mediator of the functional rearrangements resulting from psychedelics [71]. Furthermore, serotonergic psychedelics have been shown to enhance brain complexity [72], known to be decreased during states of diminished awareness, such as in DoC [73]. In addition to serotonin, we observed that increased NET transporter levels promoted recovery, which aligns with earlier research showing improved outcomes in DoC patients after administration of drugs targeting these receptors [74]. In contrast, a lower presence of dopaminergic systems was found to enhance recovery. The dopamine transporter (DAT) plays a critical role in regulating extracellular dopamine concentrations by facilitating reuptake into the presynaptic neuron. Interestingly, inhibiting this transporter has shown significant promise in enhancing arousal, as evidenced by previous studies and aligning with our results [75–78]. However, we also found that lower levels of D1 and D2 receptors correlate with increased recovery potential, which disagrees with existing literature that suggests dopamine receptor agonists improve consciousness [78]. Overall, the significant correlations between recovery profiles and neurotransmitter systems could suggest that *Inception* can also explain the long-term effects of the pharmacological interventions in DoC.

An additional feature of the *Inception* framework is its ability to model two-step interventions, which could hold considerable promise in clinical applications. For example, an initial pharmacological or therapeutical intervention could be employed to prime the brain, placing it in a more responsive and perturbable state, thereby enhancing the efficacy of a subsequent intervention, such as electrical stimulation [79]. Emerging evidence supports the efficacy of such two-step therapies, including findings that administering LSD prior to brain stimulation produces larger and distinct effects compared to stimulation alone in rodent models [80], as well as improved outcomes in depression treatment when mindfulness-based cognitive therapy precedes pharmacological interventions [81].

In summary, we introduced *Inception*, a novel *in silico* perturbation-based whole-brain modelling approach that offers subject-specific analysis and captures the perturbation’s post-acute and long-term effects. This approach improves the characterisation of perturbations needed to achieve a transition to a recovered healthy state compared to conventional methods. However, there are limitations to consider. Like previous studies, we used a structural connectivity matrix derived from healthy participants to build the DoC models. While this provides a reasonable baseline, given the heterogeneous lesions in DoC patients, it introduces potential biases that were partially mitigated by optimizing effective connectivity in each model. Additionally, the use of a parcellation with a limited number of nodes may hinder the specificity of our findings, suggesting that future research could benefit from higher-resolution atlases. Furthermore, the sample size in this study is limited, which may affect the generalizability of the results, highlighting the need for future research with larger, more diverse populations. As a next step, we plan to explore network-level perturbations rather than targeting individual nodes.

## 4 Methods

### 4.1 Participants

This study includes data from a cohort of 13 healthy controls (7 females; mean age ± s.d., 42.54 ± 13.64 years), 11 MCS subjects (5 females; mean age ± s.d., 47.25 ± 20.76 years), and 10 UWS subjects (4 females; mean age ± s.d., 39.25 ± 16.30 years). Each participant’s consciousness level was assessed by trained medical professionals using the Coma Recovery Scale-Revised (CRS-R) scale [82]. Participants diagnosed with MCS exhibited symptoms of awareness such as visual tracking, reaction to pain, or consistent response to commands. In contrast, individuals diagnosed with UWS showed arousal, eyes opened, without clear indications of awareness. This study was approved by the local ethics committee, Comité de Protection des Personnes Ile de France 1 (Paris, France), under the designation ‘Recherche en soins courants’ (NEURODOC protocol, no. 2013-A01385-40). Informed consent was obtained from the relatives of the patients. All investigations adhered strictly to the principles outlined in the Declaration of Helsinki and complied with the regulations of France.

### 4.2 Resting-state fMRI acquisition parameters

Resting-state fMRI data was collected using a 3T General Electric Signa System. T2*-weighted whole-brain resting-state images were captured through a gradient-echo EPI sequence in axial orientation, producing 200 volumes across 48 slices with a slice thickness of 3 mm, a TR/TE of 2400 ms/30 ms, voxel size of 3.4375 × 3.4375 × 3.4375 mm, flip angle of 90°, and FOV of 220 mm^2^). Furthermore, an anatomical volume was obtained with a T1-weighted MPRAGE sequence during the same session, consisting of 154 slices with a slice thickness of 1.2 mm, a TR of 7.112 ms, TE of 3.084 ms, voxel size of 1 × 1 × 1 mm and flip angle of 15°.

### 4.3 Resting-state fMRI pre-processing

Resting-state fmri data was preprocessed following the methodology detiled in [83], using FSL software (http://fsl.fmrib.ox.ac.uk/fsl). Briefly, preprocessing was conducted using MELODIC (Multivariate Exploratory Linear Optimised Decomposition into Independent Components) [84]. This process included discarding the first five volumes, correcting for motion with MCFLIRT [85], extracting the brain using BET (Brain Extraction Tool) [86], applying spatial smoothing with a 5 mm FWHM Gaussian kernel, performing rigid-body registration, high-pass filtering with a 0.01 Hz cutoff, and conducting single-session ICA (Independent Component Analysis) with automatic dimensionality estimation. Next, lesion-driven artifacts, for patients, and noise components were independently identified and regressed out for each subject using FIX (FMRIB’s ICA-based X-noiseifier) [87]. Finally, using FSL tools, the images were coregistered and the time-series were extracted from the AAL atlas for each subject in MNI space, including 90 cortical and subcortical non-cerebellar brain areas [88].

### 4.4 Anatomical data acquisition

For anatomical connectivity, we used diffusion MRI (dMRI) data from 16 healthy right-handed participants (5 women, mean age: 24.8 ± 2.5) collected at Aarhus University, Denmark. The study protocol received approval from the Research Ethics Committee of the Central Denmark Region and the internal research board at CFIN, Aarhus University. Written informed consent was obtained from all participants.

Imaging data were captured during a single session on a 3T Siemens Skyra scanner at CFIN, Aarhus University. The structural MRI T1 scan parameters included a voxel size of 1 mm^3^, a reconstructed matrix size of 256 × 256, an echo time (TE) of 3.8 ms, and a repetition time (TR) of 2300 ms.

For dMRI data collection, the parameters were as follows: TR of 9000 ms, TE of 84 ms, flip angle of 90 degrees, reconstructed matrix size of 106 × 106, voxel size of 1.98 × 1.98 mm with a slice thickness of 2 mm, and a bandwidth of 1745 Hz/Px. The dataset included 62 optimal nonlinear diffusion gradient directions at b = 1500 s/mm^2^, with one non-diffusion weighted image (b = 0) acquired per every 10 diffusion-weighted images. The acquisition was performed in both anterior-to-posterior and posterior-to-anterior phase encoding directions.

### 4.5 Probabilistic Tractography Analysis for Anatomical Data

We used structural connectivity data previously obtained from [89]. The structural connectivity matrix was averaged across participants through a three-step process. First, regions were defined based on the AAL template used in functional MRI data. Next, connections (edges) between nodes were estimated using probabilistic tractography within the whole-brain network. Finally, the data were averaged across participants. The FSL toolbox’s linear registration tool [85] was used to co-register the EPI image with the T1-weighted structural image, which was then aligned to the T1 template of ICBM152 in MNI space [90]. These transformations were combined and reversed to map the AAL template [88] from MNI space to the EPI native space, maintaining labelling values for accurate brain parcellations. Further details can be found in [89].

### 4.6 Neuroreceptor maps

We used 10 PET-derived whole-brain maps of neurotransmitter receptors, receptor-binding sites and transporters curated at https://github.com/netneurolab/hansen_receptors [42]. These encompass erotonin receptors (5HT1a, 5HT1b, 5HT2a, 5HT4, and 5HT6) and serotonin transporters (5HTT), dopamine receptors (D1 and D2) and transporters (DAT), and noradrenaline transporters (NET). Subsequently, we parcellated each map into 90 brain regions according to the AAL90 atlas, resulting in a 90 × 19 matrix.

### 4.7 Whole-brain modelling

We modelled the local dynamics of each brain area using a Stuart-Landau oscillator framework. This model characterizes each node as exhibiting supercritical Hopf bifurcation, a fundamental mechanism for transitioning from asynchronous noisy behaviour to oscillations [91]. Previous studies have effectively utilized whole-brain Hopf models to reproduce significant characteristics of brain dynamics observed across various methodologies, including electrophysiology [92, 93], magnetoencephalography [94], and functional magnetic resonance imaging (fMRI) [28, 95]. Mathematically, the whole-brain dynamics can be described by coupling the local dynamics of *N* Stuart-Landau oscillators using a connectivity matrix *C*, which is given by:

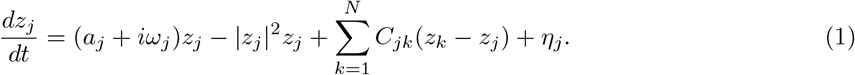

In this equation, *z*_*j*_ = *x*_*j*_ + *iy*_*j*_ represents the complex state variable of region *j. η*_*j*_ denotes additive uncorrelated Gaussian noise with variance *σ*^2^, and *a*_*j*_ represents the node’s bifurcation parameter. When *a*_*j*_ *<* 0, the local dynamics stabilize around a fixed point at *z* = 0, while *a*_*j*_ *>* 0 induces stable limit cycle oscillations with frequency 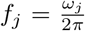. The intrinsic node frequency *ω*_*j*_ is determined empirically by band-pass filtering fMRI signals between 0.008–0.08 Hz and averaging the peak frequency of each brain region across participants.

#### 4.7.1 Linearization of the model

Previously, [89] demonstrated that setting *a* = −0.02 is the optimal operating point for fitting the dynamics of whole-brain neuroimaging. This parameter value, positioned near the bifurcation threshold, is crucial as it facilitates linearization of the dynamics, thereby enabling an analytical solution for the functional connectivity matrix *C* [96]. Employing a linear noise approximation (LNA), we can estimate the functional correlations between all brain regions. Consequently, the dynamical system involving N nodes (described in Equation 1) can be represented in a vectorized format:

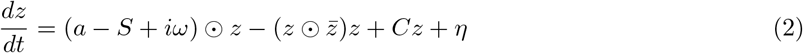

Here, **z** = [*z*_1_, …, *z*_*N*_]^*T*^, **a** = [*a*_1_, …, *a*_*N*_]^*T*^, ***ω*** = [*ω*_1_, …, *ω*_*N*_]^*T*^, ***η*** = [*η*_1_, …, *η*_*N*_]^*T*^, and **S** = [*S*_1_, …, *S*_*N*_]^*T*^, where **S** is a vector indicating the strength of each node, with *S*_*i*_ = _*j*_ *C*_*ij*_. The transpose operation is denoted by []^*T*^, the ⊙ symbol represents the Hadamard element-wise product, and 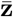 is the complex conjugate of **z**. This equation describes the linear fluctuations around the fixed point **z** = 0, which is the solution of 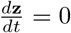. The evolution of the linear fluctuations can be described using a Langevin stochastic linear equation by separating the real and imaginary components of the state variables and neglecting higher-order terms 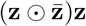:

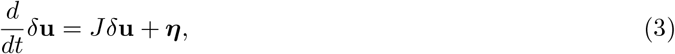

where the 2*N*-dimensional vector *δ***u** = [*δ***x**, *δ***y**]^*T*^ = [*δx*_1_, …, *δx*_*N*_, *δy*_1_, …, *δy*_*N*_]^*T*^ includes the fluctuations in both the real and imaginary parts of the state variables. The 2*N* × 2*N* matrix *J* stands for the Jacobian of the system computed at the steady state and can be represented in the form of a block matrix:

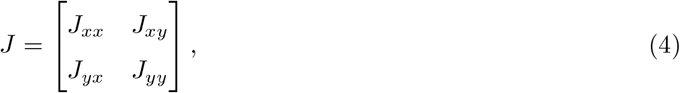

with *J*_*xx*_, *J*_*xy*_, *J*_*yx*_, and *J*_*yy*_ being matrices of size *N* × *N*. Specifically, *J*_*xx*_ = *J*_*yy*_ = diag(**a** − **S**) + *C* and *J*_*xy*_ = −*J*_*yx*_ = diag(***ω***), where diag(**v**) represents a diagonal matrix with the vector **v** on its diagonal. Importantly, this linearization is only applicable if **z** = 0 is a stable solution, meaning that all eigenvalues of *J* have a negative real part. To compute the covariance matrix *K* = ⟨*δ***u***δ***u**^*T*^ ⟩, we can express Equation 3 as *dδ***u** = *Jδ***u** *dt*+*d***W**, where *d***W** is a 2*N*-dimensional Wiener process with covariance ⟨*d***W***d***W**^*T*^ ⟩ = *Q dt*. Here, *Q* is the covariance matrix of the noise, which is diagonal if the noise is uncorrelated. Then, using Itô’s stochastic calculus, we obtain *d*(*δ***u***δ***u**^*T*^) = *d*(*δ***u**)*δ***u**^*T*^ + *δ***u** *d*(*δ***u**^*T*^) + *d*(*δ***u**)*d*(*δ***u**^*T*^). Considering that ⟨*δ***u***d***W**^*T*^ ⟩ = 0, we maintain the terms to first order in the differential *dt* and take the expectations, leading to the following expression:

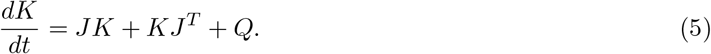

This allows us to compute the stationary covariances (for which 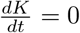) by

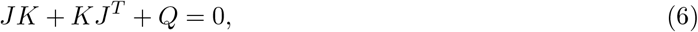

which is a Lyapunov equation that can be solved using the eigen-decomposition of the Jacobian matrix *J* [97] and, from the first *N* rows and columns of the covariance *K* we finally get the functional connectivity simulated by the model (*FC*^*simulated*^), which represents the real part of the dynamics.

#### 4.7.2 Optimization of the model

To fit the model to empirical BOLD fMRI data from each participant in each brain state, we used a pseudo-gradient optimization procedure to refine the coupling connectivity matrix *C*. This optimized matrix reflects the effective connectivity values for each anatomical connection, surpassing the dMRI-based fiber density estimations. Specifically, we iteratively compared the model’s output to the empirical functional correlation matrix (FC^empirical^), computed as the normalized covariance matrix of the functional neuroimaging data, and to the empirical time-shifted covariances (FS^empirical^(*τ*)), where *τ* denotes the time lag. These covariance matrices are obtained by shifting the empirical covariance matrix (KS^empirical^(*τ*)) and normalizing each pair (*i, j*) by 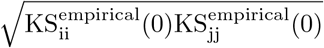. We select *τ* = 1, which is the value that minimizes the average autocorrelation. Interestingly, adjusting the time-shifted covariances can introduce asymmetries in the coupling matrix *C*.

The fitting is performed using a heuristic pseudo-gradient algorithm, iterating by updating *C* until achieving an optimal fit:

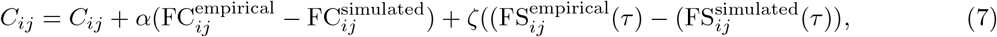

where 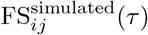 is computed by selecting the first *N* rows and columns of the simulated time-shifted covariances KS^simulated^(*τ*) and normalizing it by dividing each pair of nodes (*i, j*) by 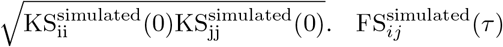 represents the simulated time-shifted covariance matrix and can be calculated as:

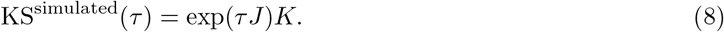

The fitting of the model is iteratively executed until a the updated value of *C* reaches a stable value, indicating that the model is optimized. *C* is initialized using the anatomical connectivity dMRI data obtained through probabilistic tractography (see Methods section 4.5), and only updated if there are existing connections in this matrix. This is followed strictly except in the homologous connections between the same regions in both hemispheres, which are also updated, as tractography is less accurate when incorporating this kind of connectivity. Moreover, *α* and *ζ* are set to 0.00001. For each iteration, we calculate the model results by averaging over multiple simulations corresponding to each participant. Finally, we refer to the optimized *C* as Effective Connectivity (EC) [98].

### 4.8 Perturbability Analysis

The initial step in our study involved defining the initial and target states. The initial state represented the brain dynamics of each MCS and UWS subject, while the target state corresponded to the brain dynamics of healthy controls.

For each MCS and UWS subject, we constructed a whole-brain model to obtain their simulated functional connectivity (FC^simulated^), their simulated time-shifted covariance matrix (FS^simulated^, hereafter referred to as the covariance matrix, COV), and their effective connectivity (EC). To define the target state, we constructed whole-brain models for each healthy subject and averaged their individual ECs, yielding the target EC.

#### *4*.*8*.*1 Inception* framework

Subsequently, for each UWS and MCS subject, we constructed a whole-brain model to apply the *in silico* perturbations. Perturbations were defined as changes in the Gaussian noise (*η*) applied to a specific node *k*. Each node was independently perturbed across a range of values from 0 to 0.5, represented mathematically as:

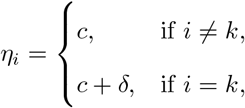

This generates perturbed FC, COV and EC matrices, capturing the short-term effects of the perturbations.

The perturbed matrices were subsequently used as inputs for a second, non-perturbed model. That is, during the optimization process, the perturbed FC will serve as the FC^empirical^ and the perturbed COV as the FS^empirical^. The output of this process includes the Inception FC, Inception COV, and Inception FC, representing the post-acute and long-term effects of the perturbation applied.

To quantify the similarity to the target group (i.e., the recovery) induced by these perturbations, we constructed the Recovery Perturbability Profile (RPP) for each subject (Figure 1b). Specifically, for each stimulated node and perturbation intensity, we computed the Frobenius norm distance between the resulting Inception EC and the target EC, yielding a matrix with dimensions *N x* perturbation values. The perturbation that yielded the lowest Frobenius norm for each node was identified as the one maximizing recovery and was included in the RPP, resulting in a vector of dimension *N*. The node with the lowest Frobenius norm in the RPP was defined as the recovery node, the corresponding RPP value was termed the recovery score, and its associated Inception EC was referred to as the recovery EC. For group-level analysis, we averaged the RPP across all subjects within the group.

#### *4*.*8*.*2 Awakening* framework

We adapted the *Awakening* framework as described in [28], which involves perturbing a whole-brain model and analyzing the resulting perturbed FC. We followed the same perturbation protocol used in the *Inception*, independently perturbing each node by varying the Gaussian noise (*η*) within a range of 0 to 0.5. The key distinction is that, unlike *Inception*, we did not construct a second model fitted on the perturbed data. Similarly, we constructed the RPP using the perturbed EC, rather than the Inception EC, and identified the recovery node, recovery score and EC for each subject.

#### 4.8.3 Comparing *Awakening* and *Inception*

The comparison between *Awakening* and *Inception* frameworks was based on the recovery scores derived from the RPP of each methodology. In addition to the RPP obtained with the ECs, we also built RPPs using the FC and COV matrices, independently. First, this required defining the target FC and COV, which we represented as the averaged FC^empirical^ and FS^empirical^ across all healthy subjects. For the *Inception* framework, we computed the Frobenius norm between the Inception FC and the target FC, as well as between the Inception COV and the target COV. For the *Awakening* framework, we computed the Frobenius norm between the perturbed FC and the target FC, and between the perturbed COV and the target COV.

### 4.9 K-means clustering analysis

We performed K-means clustering to distinguish the UWS and MCS groups on three distinct inputs: global brain connectivity (GBC), EC total degree and RPP, analyzed independently. GBC is a metric of FC, and is computed as the average correlation between each brain node’s time-series with all other nodes. The EC total degree was computed as the sum of the in-degree and the out-degree of the EC, as presented in [98]. For this analysis, we used the empirical FC and the individual EC of each subject. The algorithm was configured to identify two clusters and executed 1000 times for each input with varying initializations to mitigate sensitivity to random centroid placement. Clustering performance was assessed using accuracy and silhouette score metrics.

### 4.10 Correlation with neuroreceptor maps

To investigate potential correlations between the Recovery Perturbability Profile (RPP) and neuroreceptor maps, we independently z-scored each neuroreceptor map. Additionally, to account for significant variations in values between cortical and subcortical regions within each map, we normalized these regions separately.

To confirm that the significant correlations found were not driven by random chance we created a randomised variation of the maps preserving spatial autocorrelation. For this we used the Neuromaps library [99], specifically the moran function which applies Moran spectral randomization [100] to generate the null variations of the receptor map. We then applied min-max normalization, rescaling the data to match the original mean of the receptor map.

### 4.11 Statistical analysis

We applied a 1000-permutations t-test to compare the differences between the *Awakening* and *Inception* frameworks. For the similarity assessments to the target state, we employed a 1000-permutations two-sample t-test. The threshold for statistical significance was set to p-values *<*0.05. Additionally, we corrected for multiple comparisons using the False Discovery Rate (FDR) [101] at the 0.05 level of significance.

## 6 Supplementary material

**Figure 7.**
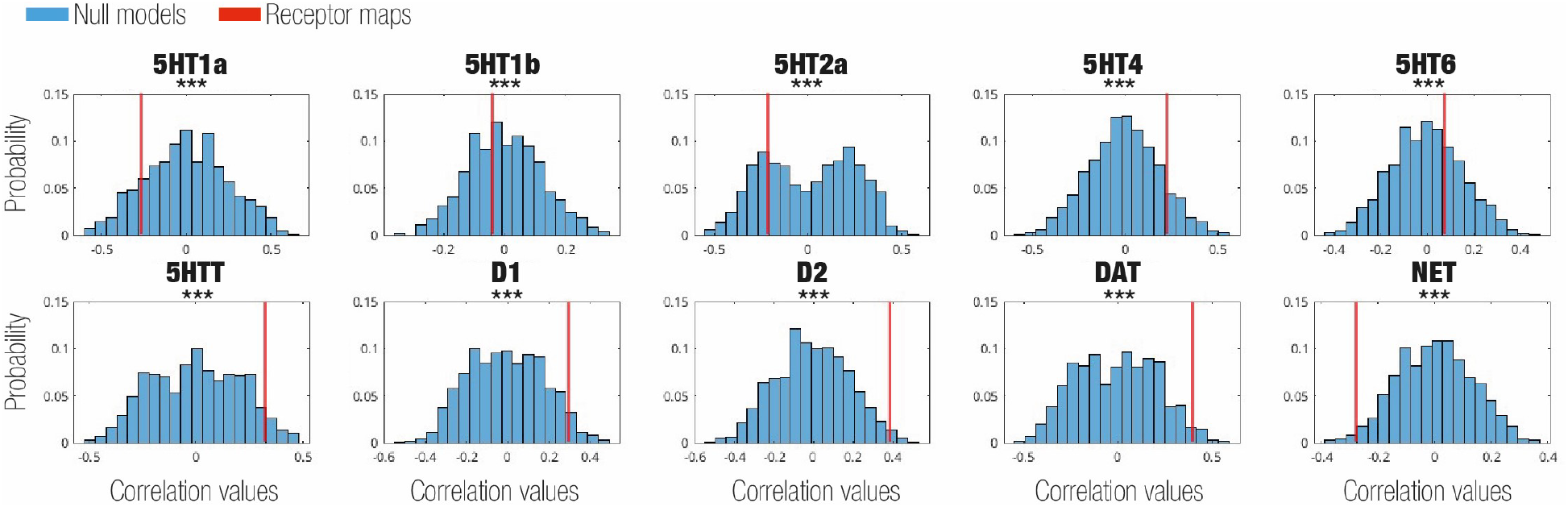
Correlation between RPP and receptor maps.

## Notes

### Competing Interest Statement

The authors have declared no competing interest.

